# Mapping Social Ecological Systems Archetypes

**DOI:** 10.1101/299693

**Authors:** Juan Rocha, Katja Malmborg, Line Gordon, Kate Brauman, Fabrice DeClerk

**Affiliations:** Stockholm Resilience Centre, Stockholm University, Kräftriket 2B, 10691 Stockholm; Beijer Institute, Swedish Royal Academy of Sciences, Lilla Frescativägen 4A, 104 05 Stockholm; Institute of Environment, University of Minnesota, St Paul, MN 55108, USA; Biodiversity International, CGIAR, Montpellier Cedex 5, France

## Abstract

While sustainable development goals are by their nature global, their achievement requires local action and thus targeting and monitoring sustainable solutions tailored to different social and ecological contexts. Ostrom stressed that there are no panaceas or universal solutions to environmental problems, and developed a social-ecological systems’ (SES) framework -a nested multi-tier set of variables- to help diagnose problems, identify complex interactions, and solutions tailored to each SES arena. The framework has been applied to over a hundred cases, typically reflecting in-depth analysis of local case studies, but with relatively small coverage in space and time. While case studies are context rich and necessary, it can be difficult to upscale their lessons to policy making realms. Here we develop a data driven method for upscaling Ostrom’s SES framework and apply it to a context where data is scarce, incomplete, but also where sustainable solutions are needed. The purpose of upscaling the framework is to create a tool that facilitates decision-making on data scarce contexts such as developing countries. We mapped SES by applying the SES framework to poverty alleviation and food security issues in the Volta River basin in Ghana and Burkina Faso. We found archetypical configurations of SES in space. Given data availability, we study their change over time, and discuss where agricultural innovations such as water reservoirs might have a stronger impact at increasing food security and therefore alleviating poverty and hunger. We conclude by outlining how the method can be used in other SES comparative studies.

---

Zero hunger and no poverty are the first two sustainable development goals (1, 2). Together with clean water and sanitation, they conform the most basic needs of human beings. Understanding how societies and ecosystems self-organise to provide these goods and services, to meet these basic needs, is a core challenge of sustainability science. Countries around the world have agreed on pursuing 17 sustainable development goals. Achieving them requires targeting and monitoring solutions that fit distinct social-ecological contexts (3). Countries must therefore understand the diversity and dynamics of social and ecological characteristics of their territories. But data to meet these demands are not always available or even collected (4), so development of methods to quantify social-ecological contexts in data scarce countries is imperative.

Nobel prize winner Elinor Ostrom advocated for embracing social-ecological complexity. Ostrom recognized that there is no universal solution to problems of destruction of overuse of natural resources (5) and further developed a Social-Ecological Systems’ (SES) framework hoping it would help accumulate knowledge and better understanding of what works and what does not in different SES contexts (6). The SES framework is a nested multi tier set of variables that has been suggested as features that characterise distinctive aspects of SES. The framework has been typically applied to local case studies that cover relatively small areas and short periods of time (documented in two publicly available datasets: the [SES Library](https://seslibrary.asu.edu/) and the [SES Meta Analysis Database](https://sesmad.dartmouth.edu/)). Over a hundred case studies have been coded in these databases. But the scale at which they are coded makes it difficult to extrapolate their lessons to arenas relevant for policy making, or to compare them to better understand what interventions work and where.

In order to target development interventions, we need to be able to characterize SES in data scarce places. The purpose of this paper is to develop a data-driven method to upscale Ostrom’s SES framework. It is targeted to countries where available data is restricted in quality and monitoring programs may not be on place. As a working example, we studied the Volta River basin, a cross national watershed that covers roughly two thirds of Burkina Faso’s and Ghana’s joint territories. The West African Sahel, where the headwaters fo the Volta are located, is a highly vulnerable area affected by wide-spread poverty, recurrent droughts and dry spells, political upheaval, emergent diseases (e.g. malaria), rapid urbanization, and growing food demand (Lambin:2014eg). The region offers a sharp gradient in climate as well as in economic development, from the relatively rich urbanized areas in southern Ghana to northern Burkina Faso, where smallholder farming and pastoralism dominates. We consider as an intervention example the construction of water reservoirs, which might have an impact on food security. The following section outlines how we operationalize the Ostrom’s SES framework to the scale of the Volta River basin and apply it using publicly available data and national statistics. Next we describe the SES archetypes found, how they change over time, and how reservoir development explain some trends. Finally, we discuss the overall results and the applicability of our methods to other data scarce contexts.

## Clustering SES

Identifying SES archetypes from data is in essence a clustering problem, this is a classification task of multiple elements by some measure of similarity. Numerous methods exist to perform clustering analysis, but before explaining the details of our choices, first we present a brief overview of what others have done when classifying SES and how our work improves previous efforts.

The idea that SES are intertwined and interdependent systems is not new: SES are human and natural coupled systems where people interact with biophysical components; they often exhibit nonlinear dynamics, reciprocal feedback loops, time lags, heterogeneity and resilience (7). It has been suggested that complex adaptive systems, such as SES, should leave statistical signatures on social and ecological data that would allow pattern identification of typologies and make it possible to follow their spatial patterns as well as trajectories through time (8, 9). Earlier efforts to map SES have been more general in purpose, and global in scale, such as the attempt to identify Anthromes (“human biomes”) (10, 11), or general land system archetypes (12–14). Reflecting on global consequences of land use, Foley *et al.* (15) proposed a conceptual framework for bundles of ecosystem services, the idea that landscape units can be classified by the sets of goods and services that a SES co-produces, or more generally, a set of social-ecological interactions. This framework has gained empirical support (16) with studies that range from the watershed to national scales in Canada (17, 18), Sweden (19, 20), Germany (21), and South Africa (22). Similar ordination methods has also been used to study regime shifts from foraging to farming societies in ancient SES (23).

Despite the differences in purpose, scale, resolution and datasets used, what the aforementioned studies have in common is that they attempt to map SES by combining multivariate methods of ordination and clustering algorithms to identify i) systems’ typologies and ii) potential underlying variables of change - what explains the variability of the typologies. However, the studies do not provide guidance on how to make choices regarding optimal number of clusters or algorithmic selection, limiting their replicability when applied to different places or data. These choices remain idiosyncratic in the SES literature and there has been no way of assessing best practices. Here we apply clustering techniques to SES data while explicitly addressing these limitations. To guide us in choice of variables, we use Ostrom’s SES framework. In Ostrom’s parlance a SES has 6 key subsystems: i) resource units (RU), resource system (RS), governance system (GS), users (U), interactions (I) and outcomes (O); all framed by social, economic and political settings (S) as well as by related ecosystems (ECO). Each of these subsystems has a nested second tier of variables (53 in total in the original proposal) aimed to capture key features of the first tier (6). We obtained publicly available datasets covering the second administrative level for Ghana and Burkina Faso (districts and provinces respectively). From these, we matched any data that in a meaningful way could be used as proxies of the Ostrom variables.

### Data and choice of variables

Since our analysis focuses on food security we use crop production as the defining key interaction (I) of our SES characterisation. It is a proxy that reflects both the capacity of the ecosystem to provide food services, and also the human input (e.g. fertilisers, labor) and preferences (e.g. crop choices) necessary to co-produce the service. Crop data, both production and cultivated area, were obtained from national statistical bureaux. While data does exist for 32 crops from 1993-2012, there are large data gaps in the time series of these data (~36%) (SM Fig 1). We thus chose 7 crops with minimum missing values (3.85%), and used averages based on the last 7 years to correct for outliers (See SM for crop selection). Users (U) and their social, economic and political settings (S) were here characterised by national census statistics and their change between the years 1996 and 2006 for Burkina Faso and between 2000 and 2010 for Ghana. The ecological system (ECO) is characterised by biophysical variables from Mitchell et al. (24) that summarises aridity, mean temperature, precipitation and slope. The resource system (RS) is a combination of variables that facilitate or inhibit crop production (our key interaction), such as the presence of water reservoirs, and the variance of food energy (kcals) produced as a proxy of predictability. Predictability is found to be associated with the capacity of self-organization and the emergence of managerial rules (6). Resource units (RU) were characterised by cattle per capita, since this is a source of insurance for farmers in the area (25). All data is standardised to the range 0:1 and log transformed for distributions with heavy tails. Table 1 summarises the framework and the proxies used.

**Table 1.**
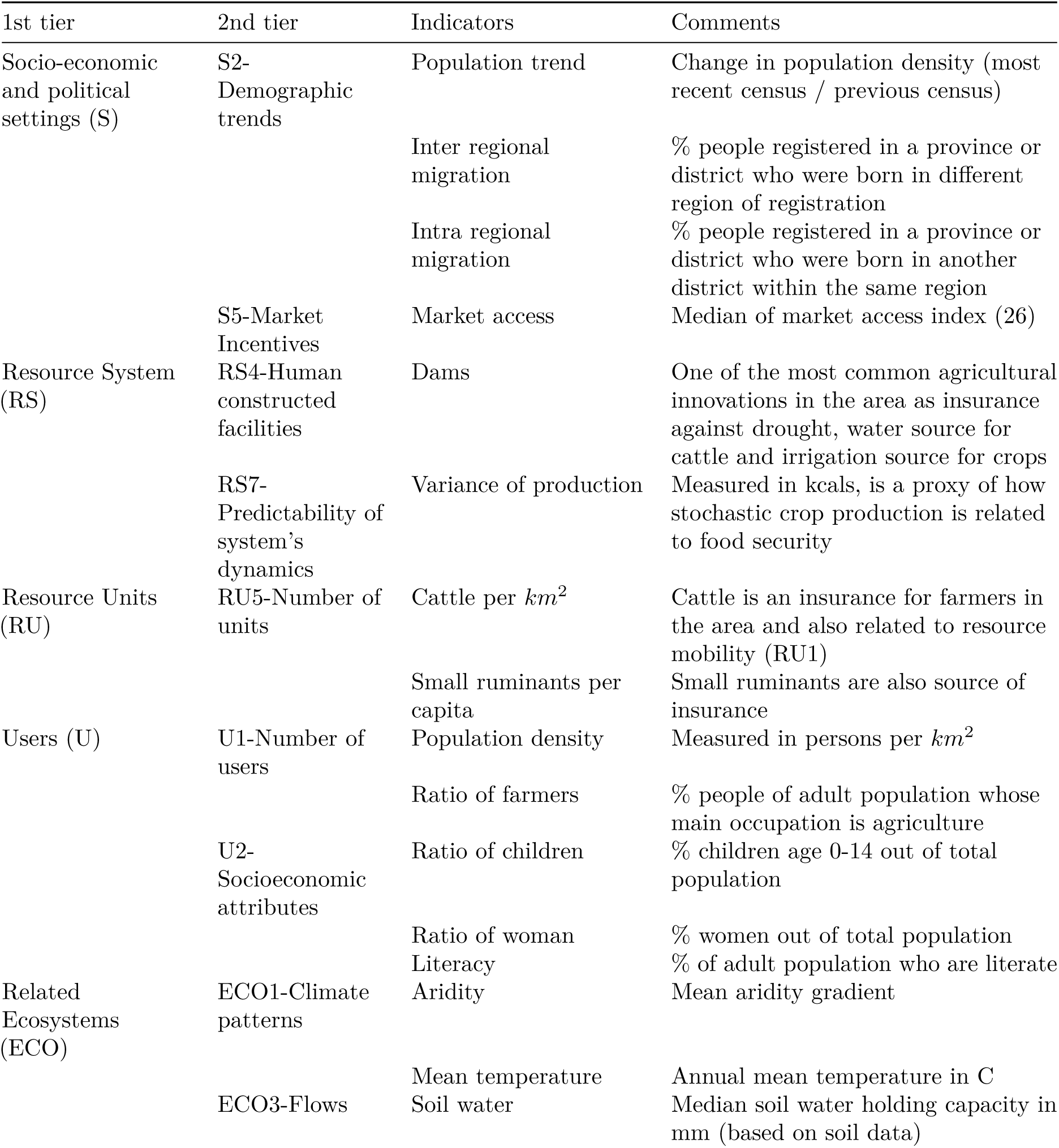
Summary variables used and their equivalence with the Ostrom’s SES framework.

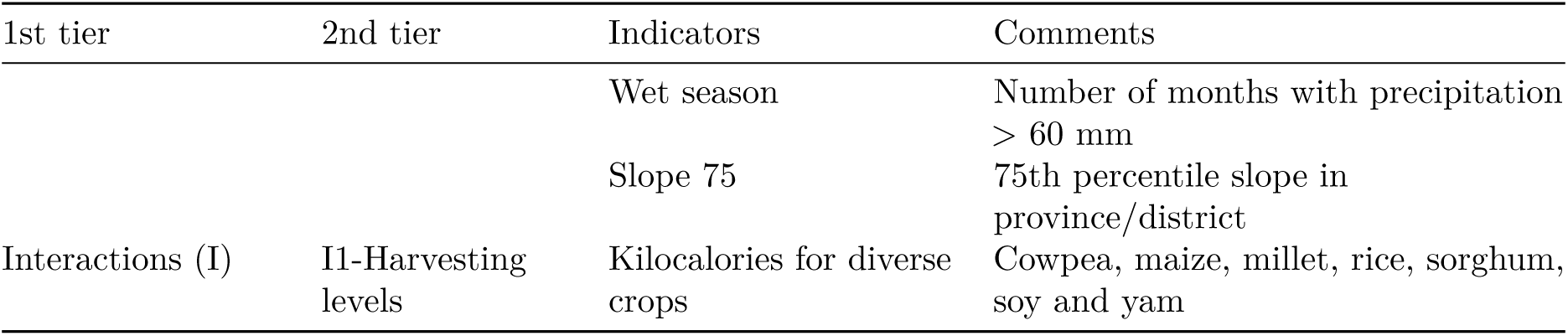

## Results

The results demonstrate that the Volta river basin is composed of distinct sets of social-ecological systems. To test the optimal number of clusters, 30 different indexes were compared following the sensitivity analysis described by Charrad et al (27) (See SM Methods). Each index is defined as an objective function (e.g. Silhouette, Duda, Dunn) that is maximised or minimised over the number of clusters to test. The clustering search identifies an optimal number of 6 archetypes suggested by 11 out of 30 indexes, followed by 3 clusters (4 indexes), and 9 clusters (4 indexes). We further test the internal validation and stability validation of 9 different clustering techniques (28). By comparing different algorithms, the test suggest that hierarchical clustering is the best performing technique with low numbers of clusters, while stability validation suggests *clara* or *pam* with higher number of clusters. Figure 1 compares the clustering of the second level administrative units into 6 clusters with the best performing clustering methods. All maps render very similar results, specially with *clara*, *pam* or *hierarchical* clustering (Fig 1??). Henceforth we favour 6 clusters as the number of archetypes given the strong support found by comparing 30 the indexes proposed (27). Subsequent results use hierarchical clustering because it outperforms *clara* and *pam* on the internal and stability validation.

**Figure 1:**
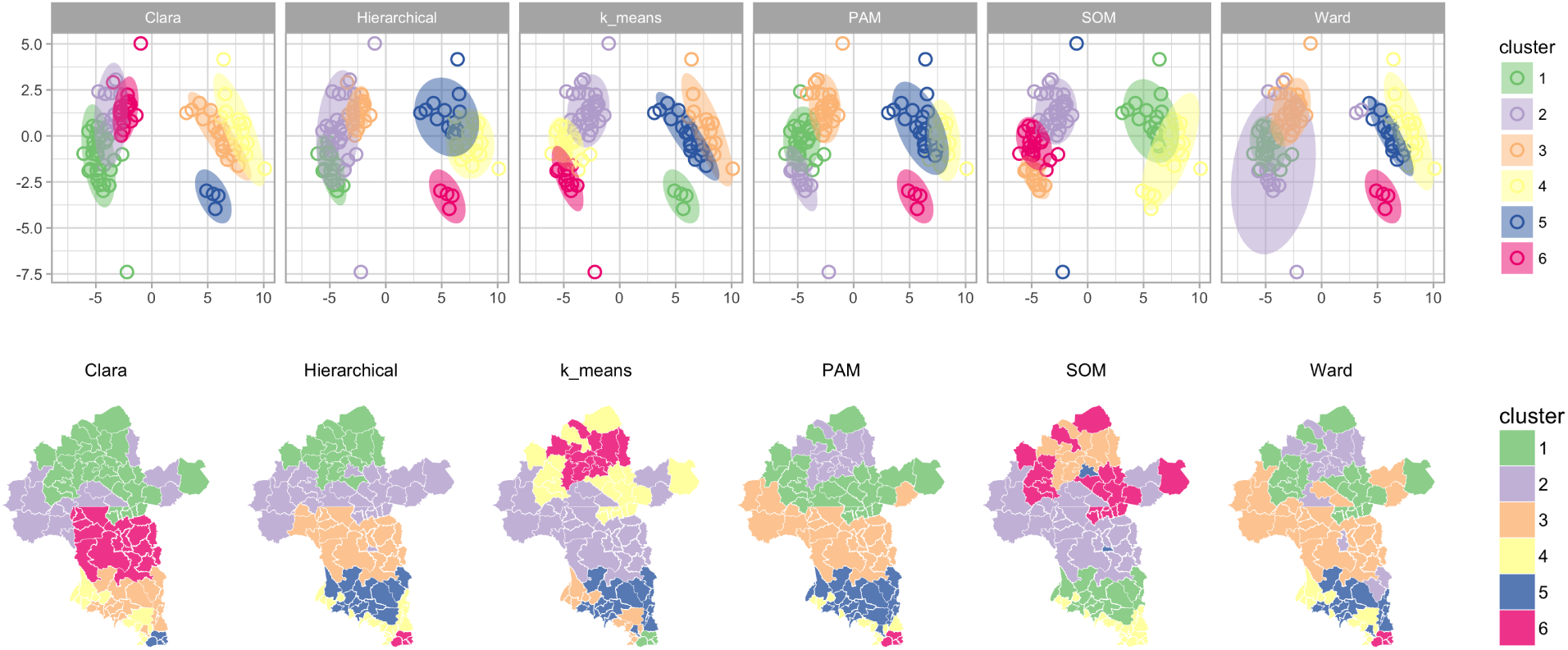
Comparison of clustering techniques. Social-Ecological systems archetypes found by applying different clustering techniques (See SM Methods for algorithm description and cluster number selection). Upper panel shows non-metric multi dimensional scaling of the data, and the lower panel the respective geographical distribution. Best performing algorithms are clara, partition around medioids, and hierarchical clustering.

Following the analogy of bundles of ecosystem services (15, 17), we also map the sets of SES variables that co-vary in space using the archetypes found by the clustering analysis (Fig. 2). As expected, SES archetypes follow a north - south gradient, with SES in the north (cluster 1 and 2) characterized by arid environments.

**Figure 2:**
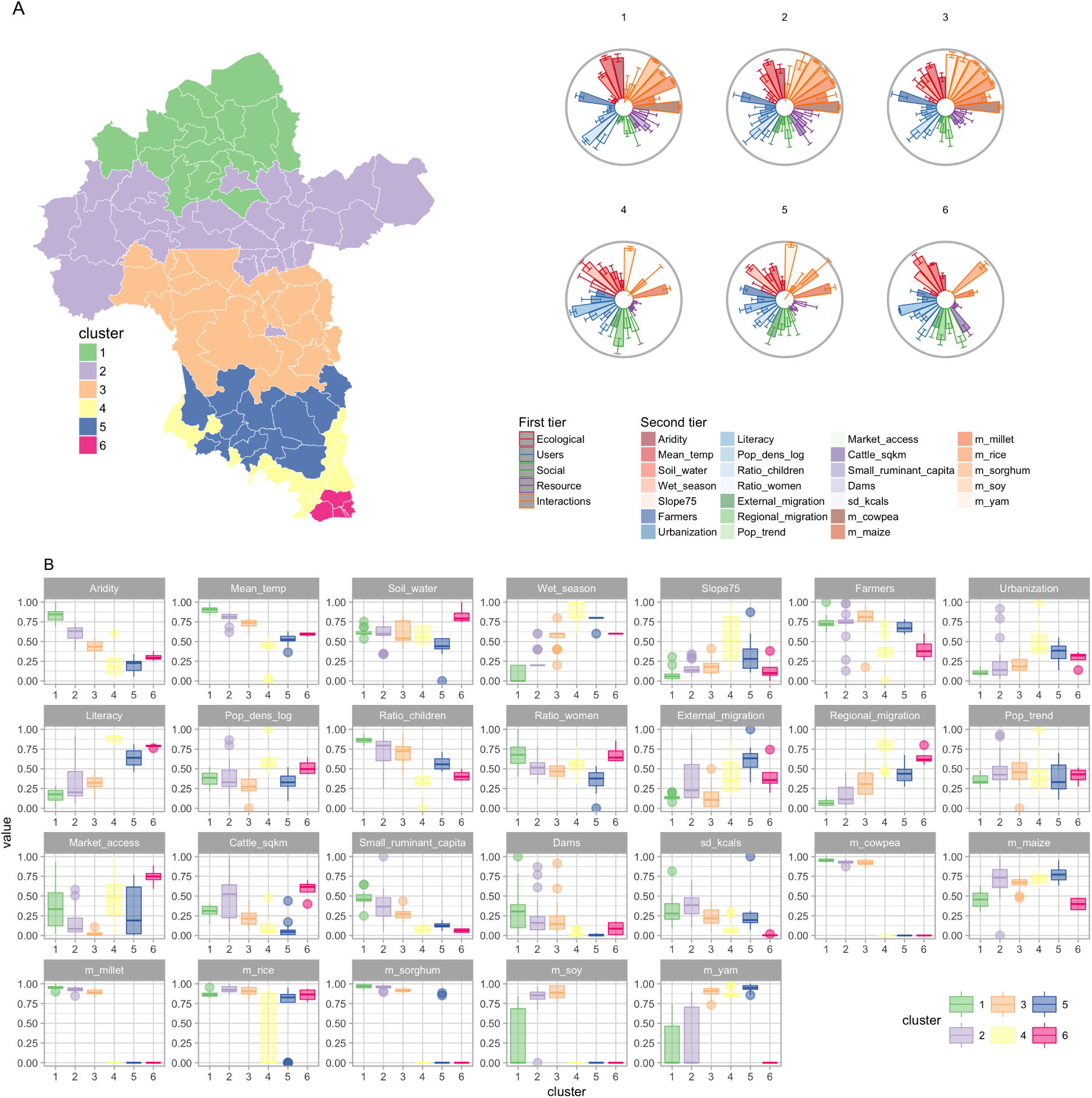
Bundles of SES variables. The SES framework was operationalized by analyzing datasets that are used as proxies forOstrom’s suggested variables at spatial units that correspond to the second administrative level (A). The clusters correspond to the best partition obtained with Hierarchical clustering (Fig 1). For each cluster there is a flower diagram of average normalized units for each second tier variable. The outer gray circle shows the unit circle, if the mean is equal to one the bar would be as high as the circle’s perimeter. The base of the bars shows the zero values. The height of the bar exhibits the variable mean for each cluster, while the error bars show standard deviations. Note that for some variables the error bars go above 1 or below zero, indicating that the mean is not necessarily the best summary statistic for all variables. (B) shows the distribution of each variable and allows for cluster comparison within variable; Supplementary Material Fig 3 complements this figure by showing the intra cluster variation while Supplementary Material Fig 4 shows the statistical differences between clusters per variable.

**Figure 3:**
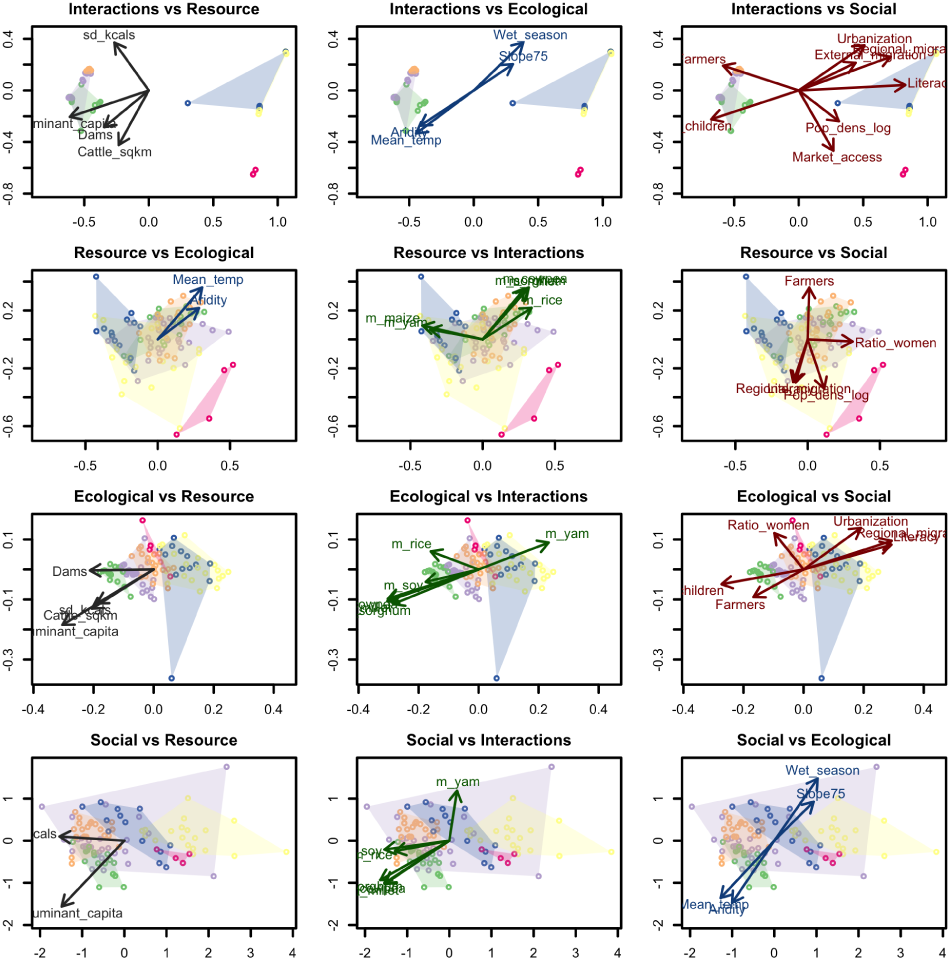
Relationships between SES framework components. Each subsystem in the Ostrom’s framework is ordered with non-metric multidimensional scaling and vectors are fitted for all other variables that significantly (p < 0.001) explain the variation of the ordination. Each plot title shows the dataset in which the ordination was applied versus the dataset used for the vector fitting. The colours of points and contours correspond to the archetypes found in Fig 2.

**Figure 4:**
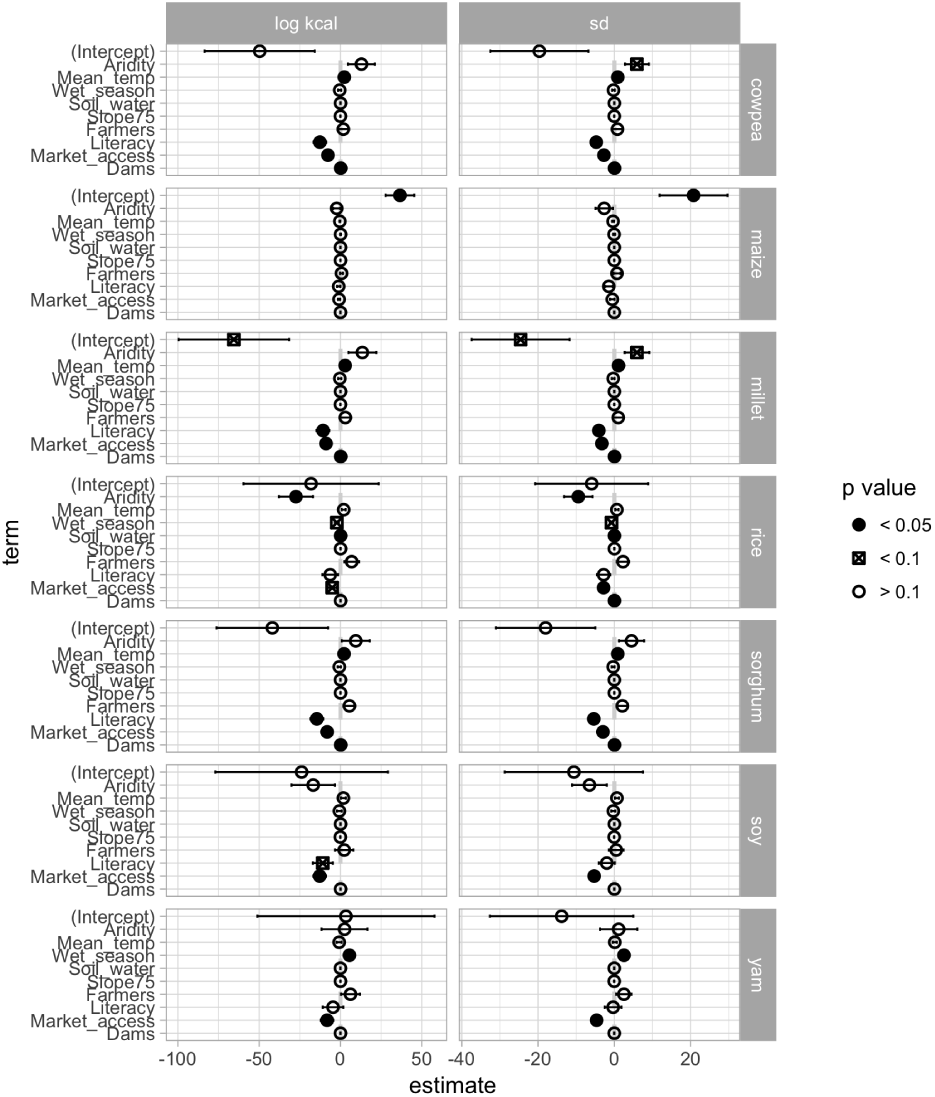
Influence of water reservoirs in kilocalorie produced per crops and its variability. The kilocalorie and its standard deviation in log scale is regressed against several factors that can influence crop production. Dams have a significant effect in cowpea, milet, rice and sorghum crops after controlling for biotic and abiotic factors that can improve production. Cropped area was controlled in the regression but excluded from the figure given that its coefficient is an order of magnitud larger, making difficult the interpretation of other coefficients. Complete regression tables are available on the supplementary material (Tables S3-S5).

In these clusters, the kilocalories produced come primarily from cowpea, millet and sorghum, with relatively higher rice and maize production in southern Burkina Faso (cluster 2). Though clusters 1 and 2 are similar, cluster 2 has in average higher cattle per *km*^2^, both higher intra- and inter-regional migration, higher literacy rates, faster population growth, but less market connectivity, and a lower ratio of women and children (SM Fig 2,3). Cluster 3 concentrates higher kilocalorie production with relatively high production of all crops analysed except for maize. Maize reaches its production peak in the south (clusters 4-5). Clusters 4 and 5 are also characterised by the production of yam and rice, lower cattle per *km*^2^, and fewer small ruminants per capita. The highest urbanisation and literacy occurs in clusters 4 and 5. External migration is low in cluster 3 but increases again further south in clusters 4-6. Cattle per *km*2 and market access reach their highest in cluster 6. The south is also dominated by longer wet seasons and higher soil water content. Note that while water reservoir density is higher in cluster 1, the variance of kilocalorie (our proxy for predictability) is high both in clusters 1, 2, 3 and 5. Later we expand on this result by testing the relationship between density of water reservoirs, kilocalorie produced and its variance with a linear regression explained below. Figure SM3 and SM4 further explore significant differences between clusters for each second tier variable.

**Figure 5:**
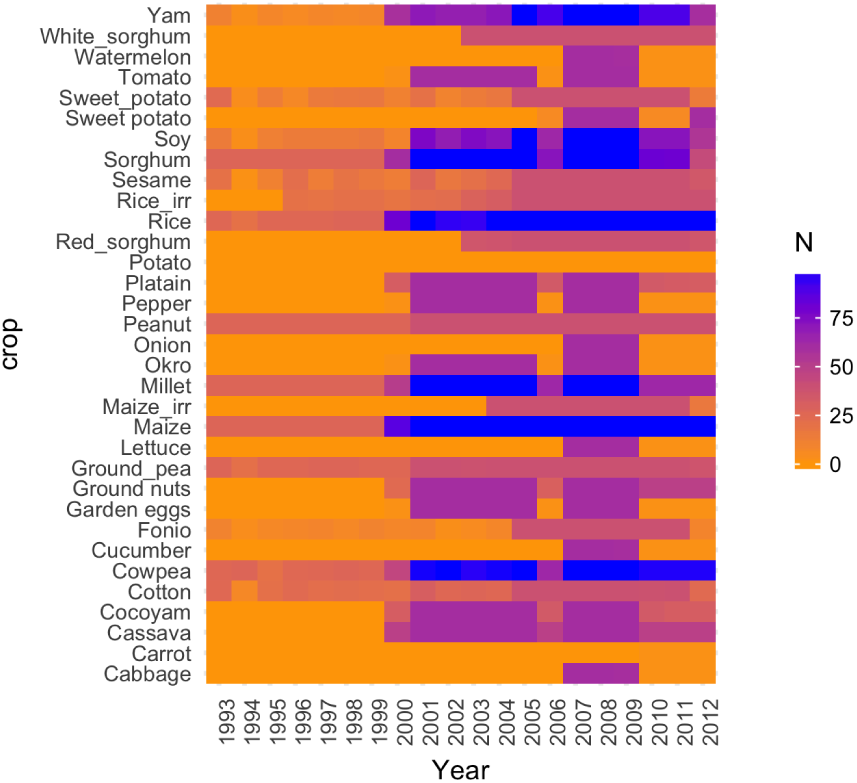
Crop selection. Number of observations (N) per crop per year. Note that there is only 7 crops with almost complete data for 7 years from the complete dataset.

**Figure 6: SM Fig1.**
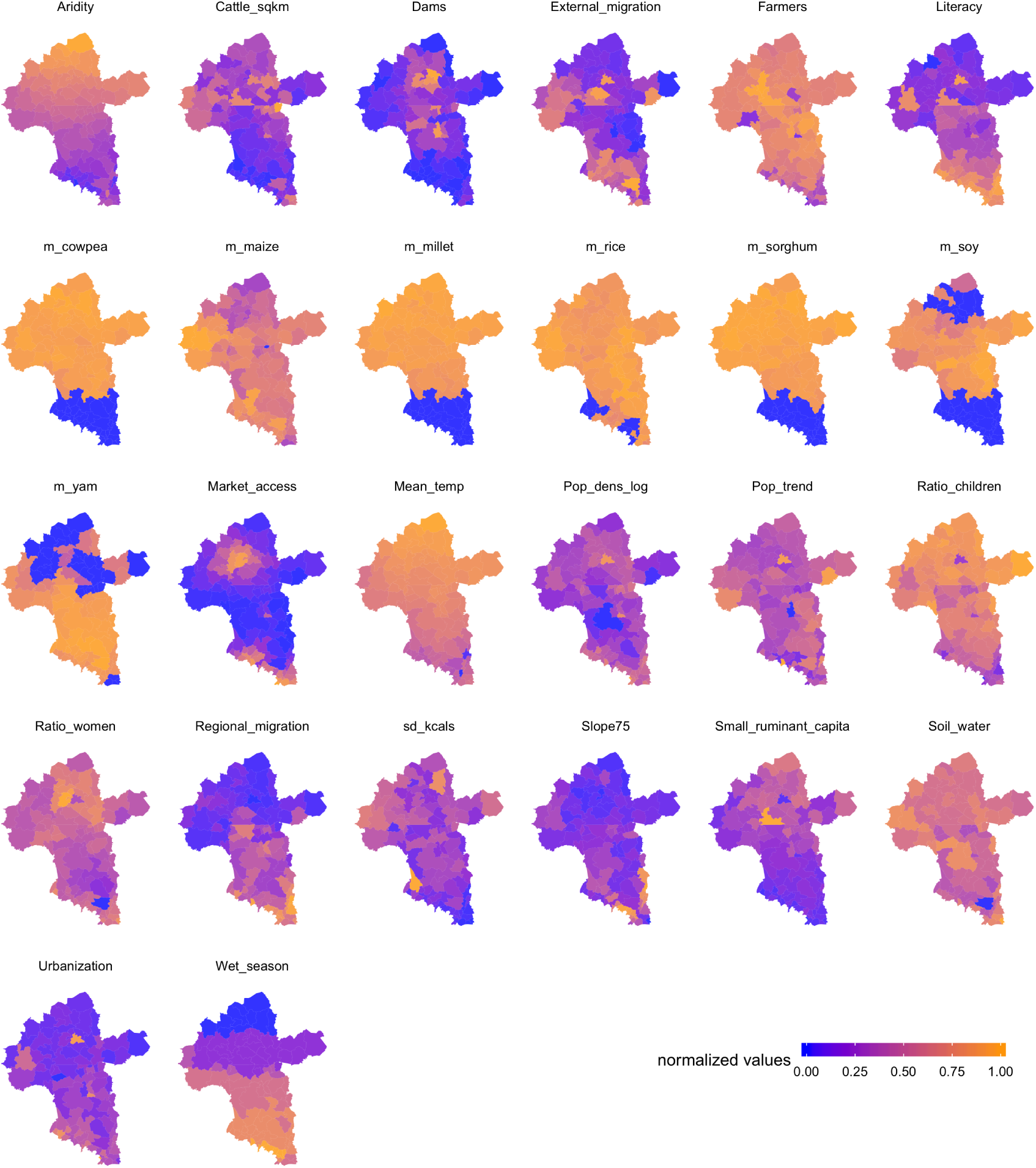
Normalized values for second tier variables and their distribution in space

**Figure 7: SM Fig2.**
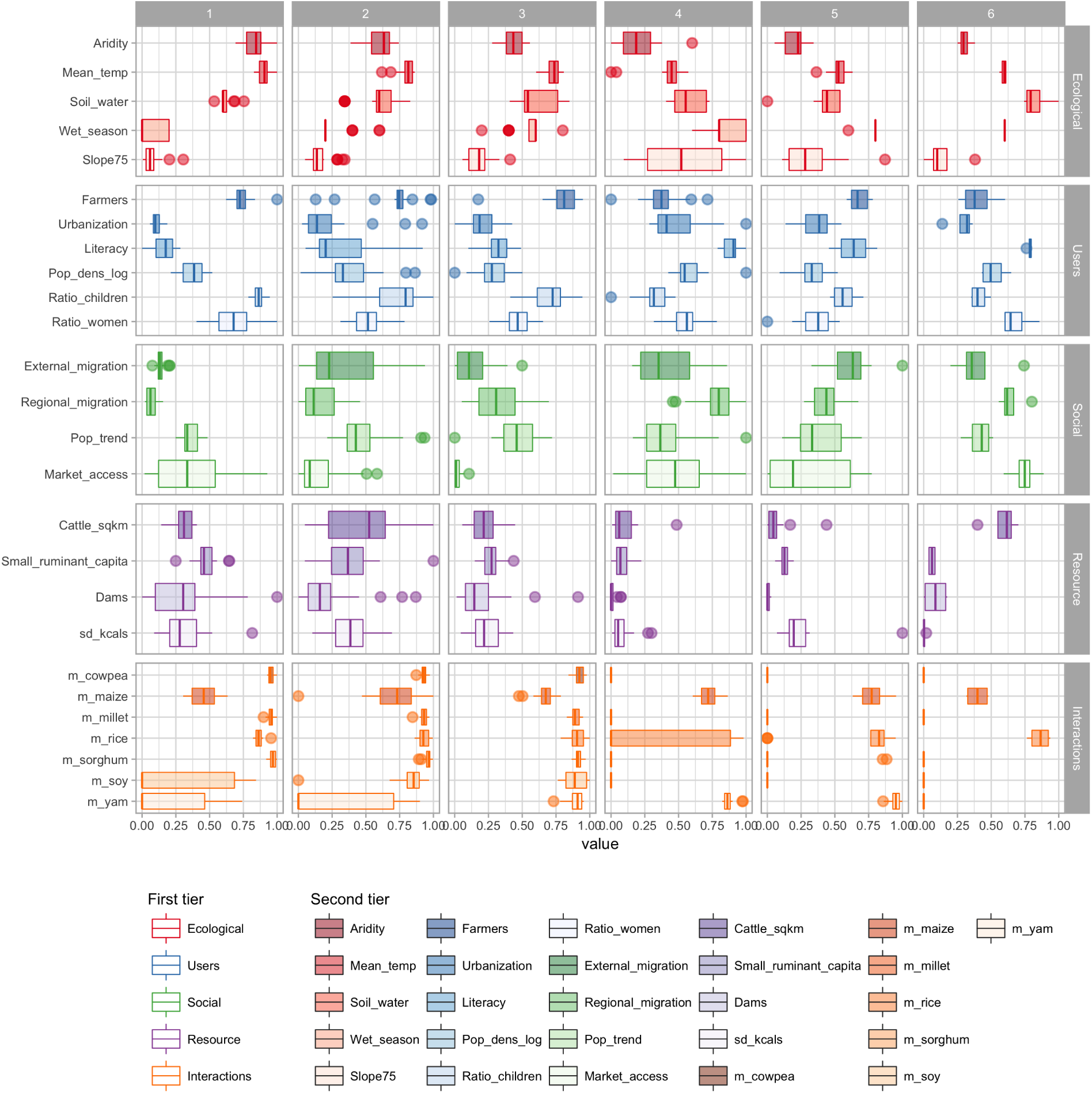
Distribution of second tier variables per cluster.

**Figure 8: SM Fig3.**
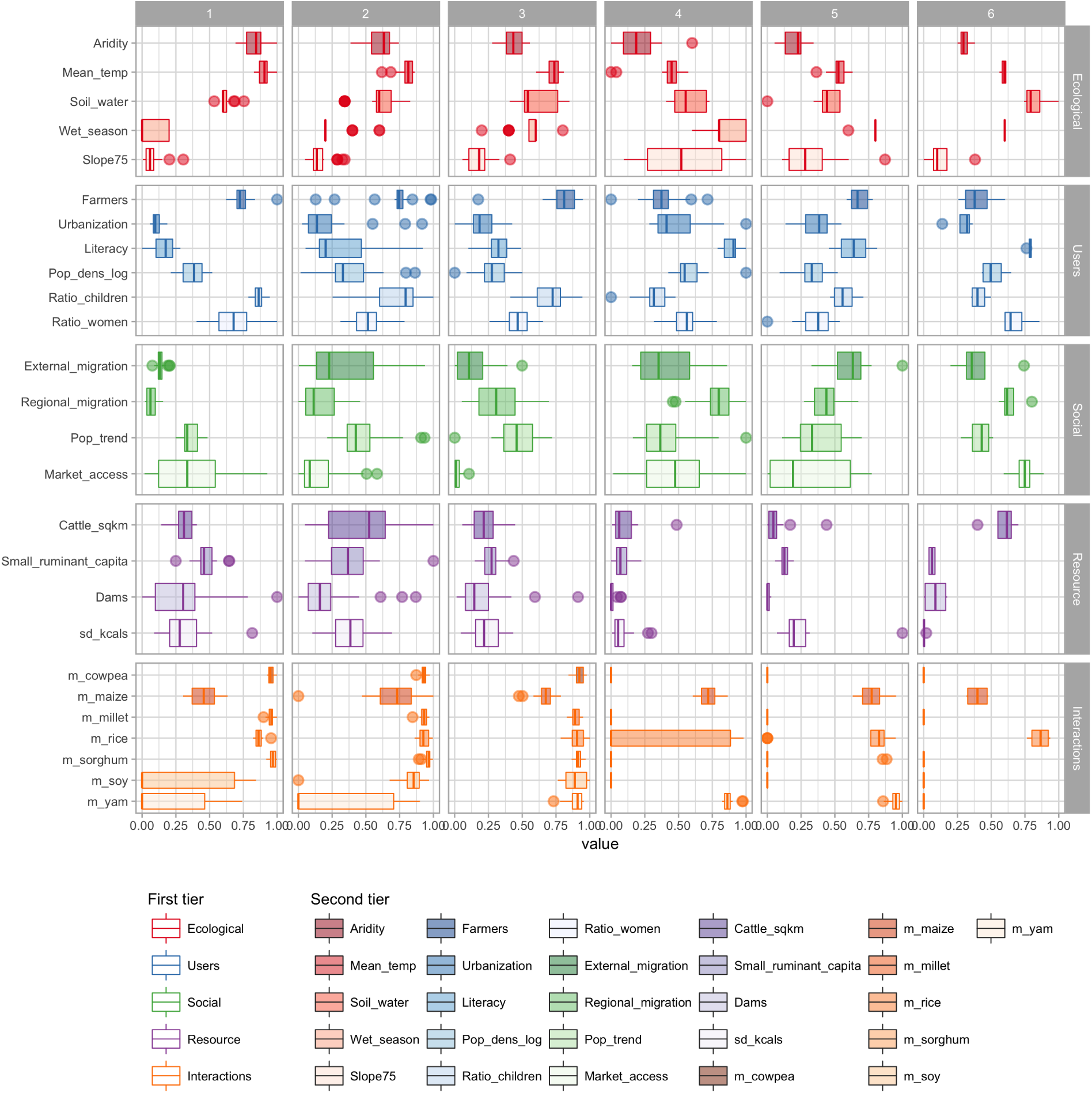
Differences between clusters per second tier variable. After an analysis of variance was performed for each variable with respect to each cluster, a Tukey test of difference is shows which clusters are significantly different or not from each other by each second tier variable analysed.

**Figure 9: SM Fig 4.**
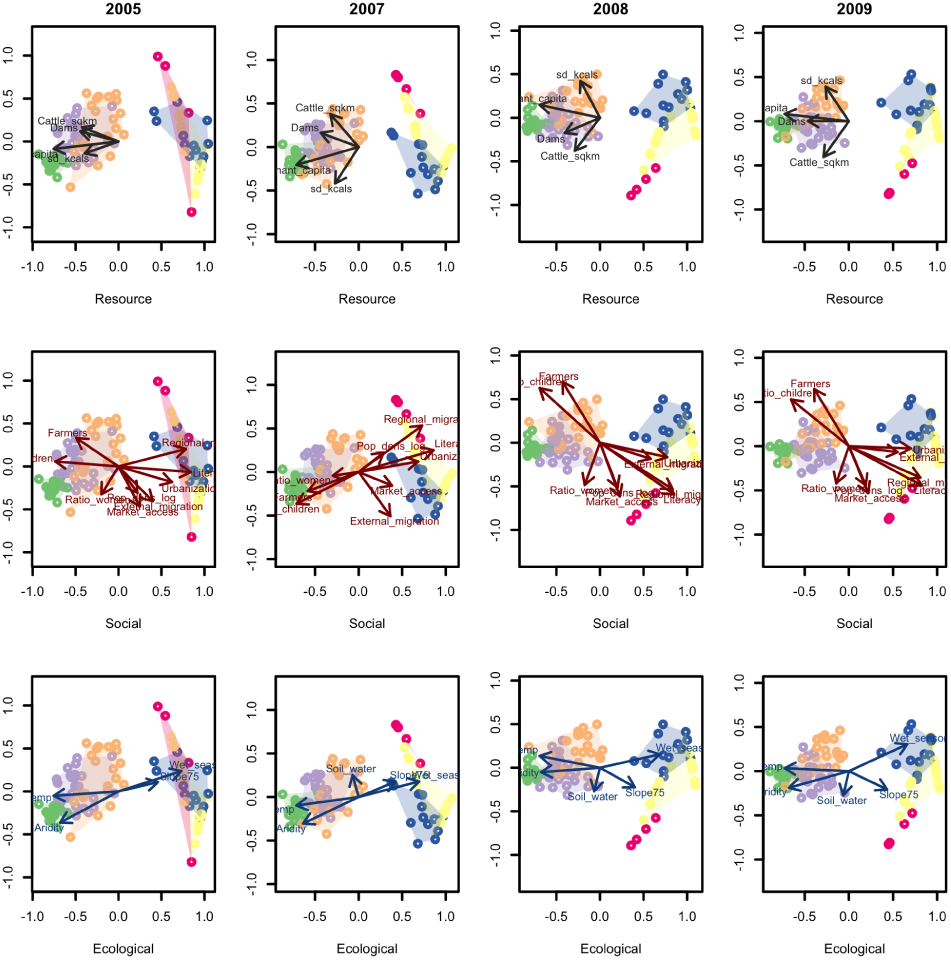
Time robustness. The interaction dataset is ordered with non-metric multidimensional scaling for multiple points in time and vectors are fitted for all other variables that significanlty (p < 0.001) explain the variation of the ordination. Each column is a different year with complete data, while each row corresponds to the dataset used for the vector fitting (resource, social, ecological). The colors of points and contours correspond to the archetypes found in Fig 2.

The potential relationships between Ostrom’s framework components were further investigated by applying vector fitting to non-metric multi-dimensional scaling (Fig 3). We use this approach to better understand what explains the variability of the archetypes found. The ordination method applied to each set of variables reveals that the clustering (Fig 2) is highly driven by the interactions (crops) data (Fig 3 top row). Clusters 1-2 tend to produce similar crops and rely heavily on cattle and small ruminants; they are also where dams are more abundant. A similar ordination on the ecological variables (ECO) supports the idea that water reservoirs and cattle have been highly correlated to places with high aridity, and occur in places where the crop portfolio is characterised by soy, cowpea, sorghum and rice. The predictability of the resource system, measured as the variance of kilocalorie production over 7 years of data, shows that clusters 2-3 are the areas where crop productivity is more unpredictable, and thus where food security might be compromised. They are also areas with higher densities of farmers and children. These relationships across the different components of the Ostrom’s framework holds when looking at years where complete crop data is available (SM Fig 4, also see SM Fig 1 for data gaps). It suggests that the results are robust across time despite limited longitudinal data.

These results put together suggest that agricultural innovations related with water reservoirs development have had more influence in northern parts of Burkina Faso (cluster 1, Fig 3). In addition, our analysis suggests that additional reservoirs would likely have a strong impact on the SES represented by clusters 2 and 3, because in these regions social and ecological conditions are similar to cluster 1 and food security is vulnerable (based on a high standard deviation of kilocalorie production). To test this hypothesis, we regressed the kilocalorie produced per crop against dam density, controlling for biophysical factors (e.g. aridity, wet season length, soil water) and social factors (literacy, number of farmers, market access) that can impact yields. Figure 4 shows that the presence of dams has a relatively small effect when compared with other coefficients, but positive and significant on the mean kilocalorie production for cowpeas, millet, and sorghum (p < 0.01), while it does have an effect on the standard deviation of the kilocalories produced for cowpea, millet, sorghum (p < 0.01) and rice (p < 0.05).

While these results confirm our speculation that dams can help increase food security in certain SES contexts, they cannot be interpreted as causal effects. Our data does not provide extensive time series, instrumental variables, or randomized control trials required to test for causality. In addition, other factors that are expected to play a role on the production of crops are not controlled for due to lack of data at the appropriate spatial scale, such as the use of fertilizers and pesticides. In summary, we can identify patterns of where the abundance of reservoirs does correlate with production of certain crops, and based on this pattern triangulate where an additional dam is likely to have a similar effect, or more importantly, where an additional reservoir is not likely to have an impact on food security. Further tests for causality would require longitudinal data on water reservoirs (when were they build, water storage capacity, location), complete longitudinal data on water and irrigation intensive crops such as vegetables, and proxies for fertilizers and pesticides use at the second-level administrative division.

## Discussion

The purpose of this paper was outlining a data driven routine for operationalising Ostrom’s SES framework and mapping SES archetypes. The method can be executed using exclusively publicly available data and open access software (27, 28), making suitable for replication in other data limited settings beyond our test case in the Volta river basin. By applying clustering with a sensitivity analysis routine to the Volta river basin case, we have demonstrated how the method performs in a setting with restricted data quality and still renders useful insights. We have found that the Volta basin can be best described by 6 SES archetypes strongly characterised by their crop productivity profiles but also by social variables such as urbanisation, literacy, and migration. Our results also suggest that the construction of water reservoirs can improve food security in places in the basin historically exposed to high variability in food security.

Our work extends previous efforts for mapping SES (17–22, 29) in that it considers a broader range of both social and ecological variables outlined by Ostrom. Our work contributes, to our knowledge, a first attempt to upscale Ostrom’s framework to a multi-national scale that matches the scale of the resource flow dynamics: the basin. Previous efforts have relied heavily in qualitative data, which restricts the analysis to smaller sample sizes (i.e. see work by (29), N=12). While our approach looses some of the richness provided by case studies, it allows us to take advantage of data over a larger spatial scale and few observations across time to draw comparisons among diverse places.

Our approach is limited by data availability. For example, we have not included any variable in the Ostrom SES framework that describes the governance of the system (G), or the use of fertilizers (RS). Although indicators of governance and fertilizer use do exist at the national scale (e.g. governance indicators, World Bank database, only available for Ghana; fertilizers FAO database), they cannot provide insight at the scale of this study, limiting our conclusions. This suggests that including governance indicators such as how often people share food, what is the structure of the social networks, or efficiency of local institutions at managing existing water infrastructure in national monitoring programs such as census or national surveys would markedly improve the ability to characterize SES. If and when these type of data become available for the Volta basin or elsewhere, they can be easily incorporated into the SES analysis here proposed.

Our work also improves replicability and reproducibility compared to previous efforts at mapping SES (17–22). Previous work has relied on only one clustering technique and relied heavily on context dependent knowledge to make subjective decisions about the number of clusters to fit and the clustering technique to apply. While this is a valid approach, it limits scalability and reproducibility because choices made for one place may not be appropriate in other places. Here we have used an updated routine with a sensibility analysis that helps the researcher to make such choices guided by the patterns already contained in the data. While a machine learning can never replace the richness of local knowledge, it does facilitates the practical application of the method in absence of in-depth qualitative data (e.g. lack of coverage), and in settings where field work validation is restricted (e.g. war zones). This approach can thus complement and guide where qualitative research efforts could be most effectively deployed. In addition, though we cannot claim causality, the patterns here presented can be useful for policy making or identifying priority areas for future investments. Qualitative and quantitative efforts at a local scale will provide ground validation of our results and inform aspects not analysed here such as governance variables.

## Conclusion

Advancing theories on sustainability science requires articulating existing SES frameworks to generalisable and replicable analysis of large scale systems. Achieving the sustainable development goals depends on distinguishing where a sustainable solution is context dependent or where it could be generalised to different SES arenas. Here we have advanced methods to identify SES by updating clustering routines with a sensitivity analysis that allow us to reinterpret a binational dataset in the Volta basin. By doing so, we identified where and under which conditions an agricultural innovation such as water infrastructure development is likely to influence the food security of farmers thriving in one of the most arid areas of the world. These patterns, although descriptive, can inform policy decisions. We believe that identifying patterns of variables in space and time that characterise different social ecological systems is key for further developing theories of sustainability, testing when interventions work, and mapping how nations progress towards sustainable development goals. The methods here outlined are generalisable to other developing country settings, and we hope they will help rigorously test under which conditions the Ostrom’s SES framework can have policy relevant implications.

## Supplementary material

### Crop data selection

Crop data records available for Ghana and Burkina Faso go as back as 1993. Yet, out of the 32 crops recorded, complete data for the basin is only available for 7 crops used in the current analysis. The dataset is far from complete. On its most recent version, the data has 31200 observations. An observation is a datapoint of a crop in a district with some production (tons) and some cropped area (ha). Out of the 31200 observations 11449 are missing values: `NA’ s` or empty cells on the raw data. This is over 36% of the most complete version of the dataset. For the analysis presented in the paper we only used the 7 most complete crops, and still we had missing values. SM Fig1 shows the missing values in orange, the darker the blue the better, meaning data is complete for the 99 spatial units analyzed. All data was log-transformed, total crop area calculated as well as the proportion of cultivated area per crop. However area values were not used for the analysis due to high correlations with production data. Production in Tons was transformed to Kcals with data from FAO. And the mean for a 7 year period (skipping 2006) will be used for the interaction dataset in the clustering analysis. The data for seven key crops has 4851 observations of which 4664 does not contain missing values. Now instead of >30%, missing values were reduced to < 4%. Mean values and log-transformations were performed for each district or province dropping the missing values, meaning that for some provices and some crops the mean is not based on 7 values but fewer (>4 in minimum cases). In doing so we avoid zero inflation on the distributions of the original data that can bias the clustering results.

### Methods

The optimal number of clusters was tested by comparing 30 different indices and their performance following a sensitivity analysis. In general, each index requires maximisation or minimisation of some measure, that helps the researcher decide the optimal number of clusters in the data. A description and interpretation of each index can be found in Charrad et al (27). We further test the internal validation and stability validation of 9 different clustering techniques: hierarchical clustering, self-organizing maps, k-means, partitioning around medoids *pam*, divisible hierarchical algorithm *diana*, a sampling based clustering *clara*, a fuzzy clustering *fanny*, self-organizing trees *sota*, and model-based algorithm (28). While the first technique offers a robust estimation of the number of clusters in the data, the second helps choosing an optimal clustering algorithm. Results are typically presented as non-metric multi dimensional ordinations and their spacial distribution in maps of the Volta basin. Alternatively, principal component ordination results are shown in Supplementary Material. The maximum dissimilarity distance was used to maximise the distance between components, while the Ward aggregation method was used to minimise the total within-cluster variance (27). For visualisations we used a less restrictive Manhattan distance to ensure convergence. We further investigate the interdependences between Ostrom’s nested variables, by reiterating the ordination on a set of variables of interest (e.g. interactions [I]) and performing vector fitting with the remaining variable sets (e.g. resource (RU and RS), social (U and S), ecological (ECO)). The same procedure was performed over time for year with complete data to test how robust are our estimates over time.

#### Principal component analysis

An alternative approach to ordination is PCA. As we can see from the ordination plots the first principal component explains ~45% of the variability in the data while the second ~12%.

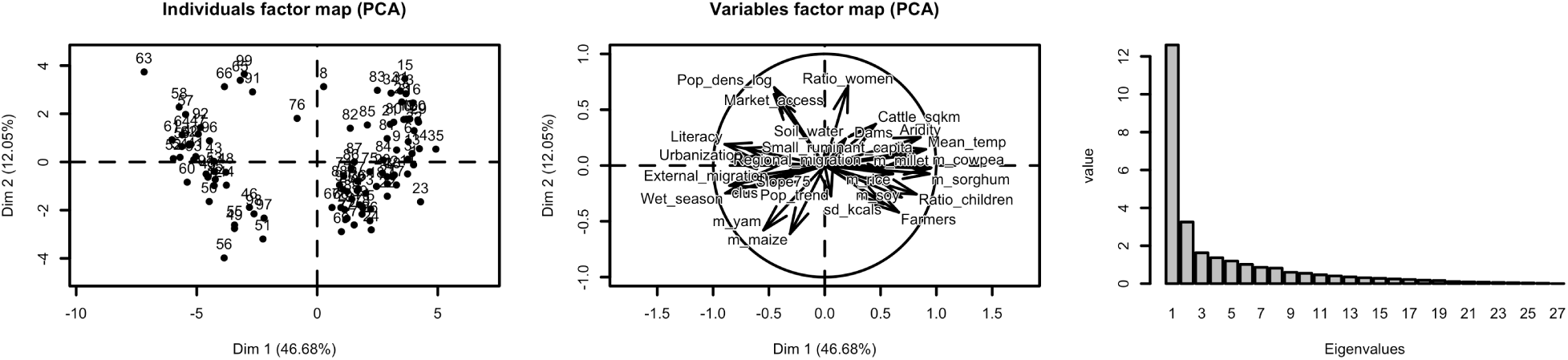

Similarly to the results presented with non-metric multidimensional scaling (Fig 1), PCA separate two big groups of clusters across the first component with a lot of overlap across the second component. The vectors confirm that clusters 1-3 are dominated by agroecosystems whose production focuses on millet, cowpeas, rice and sorghum. Clusters 1-2 have the higher aridity and mean temperatures, but also the higher concentration of water reservoirs and cattle per square kilometer. Cluster 3 has the fasters growing population and also the highest variability in crop production as well as the highest concentration of farmers. Clusters 4-5 in the south tend to be more urbanised, have higher rates of literacy, higher migration (both internal and regional), as well as agroecosystems dominated by the production of yam and maize. Cluster 6 has particularly high population density and market access.

SM Table 1 shows the loadings of each component. On the first component, variables with high positive values such as aridity, mean temperature, and produced kilocalores of cowpeas, millet or sorghum separate apart clusters 1-3; while variables with negative values such as wet season, literacy, regional migration and urbanization are responsible for the differentiation of the clusters 4-6. On the vertical axis (the second component) variables responsible for explaining variation includes population density, ratio of women, market access and production in kilocalories of yam and maize.

**Figure 10:**
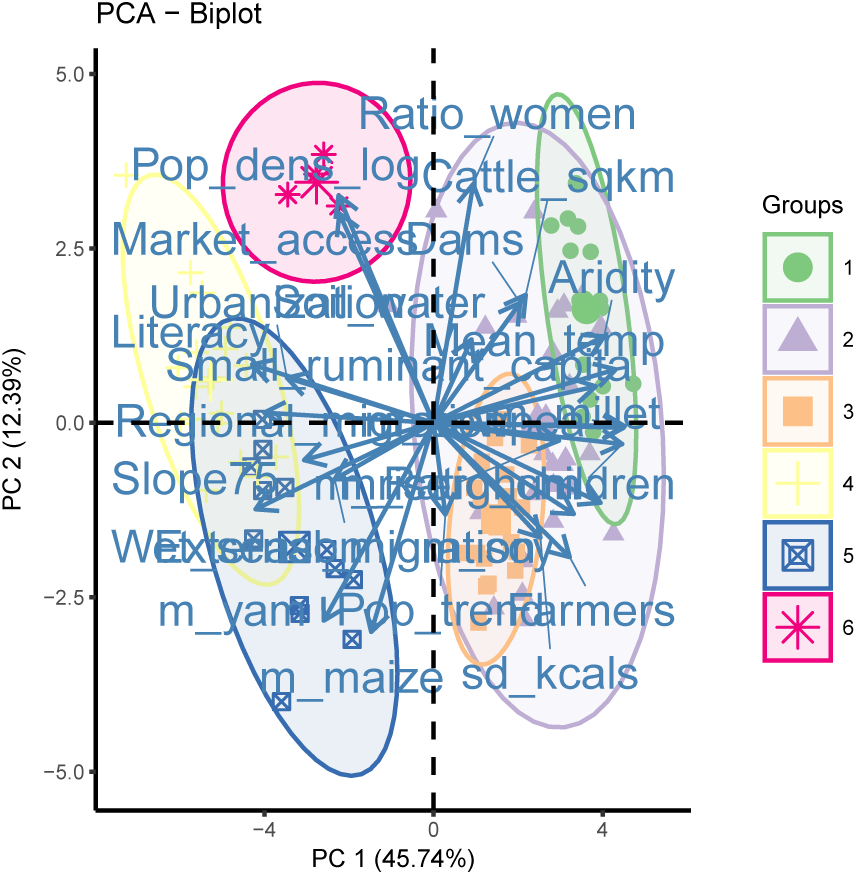
Graphical illustration of differences in variables between archetypes based on PCA.

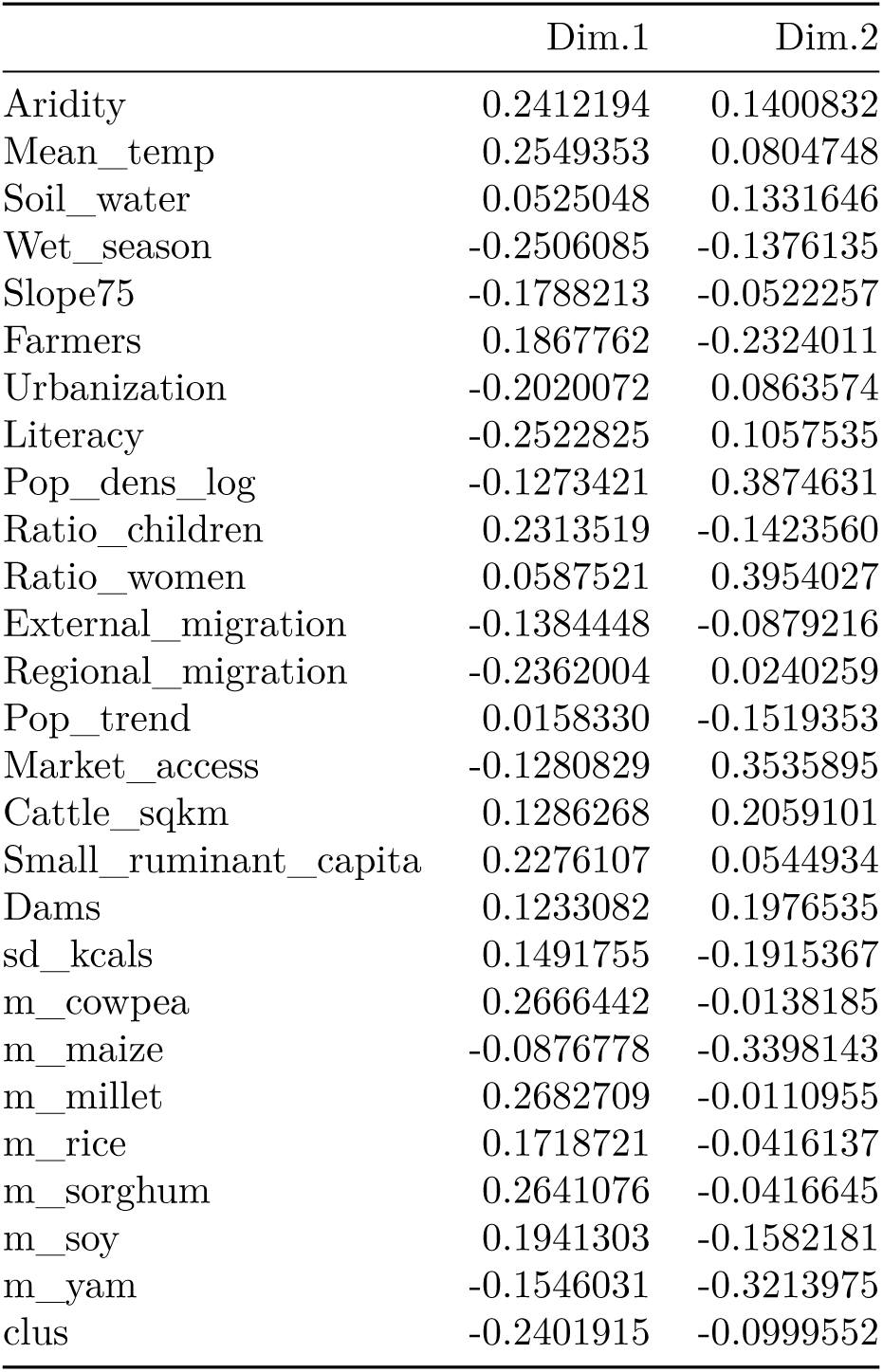

**Table 3:**
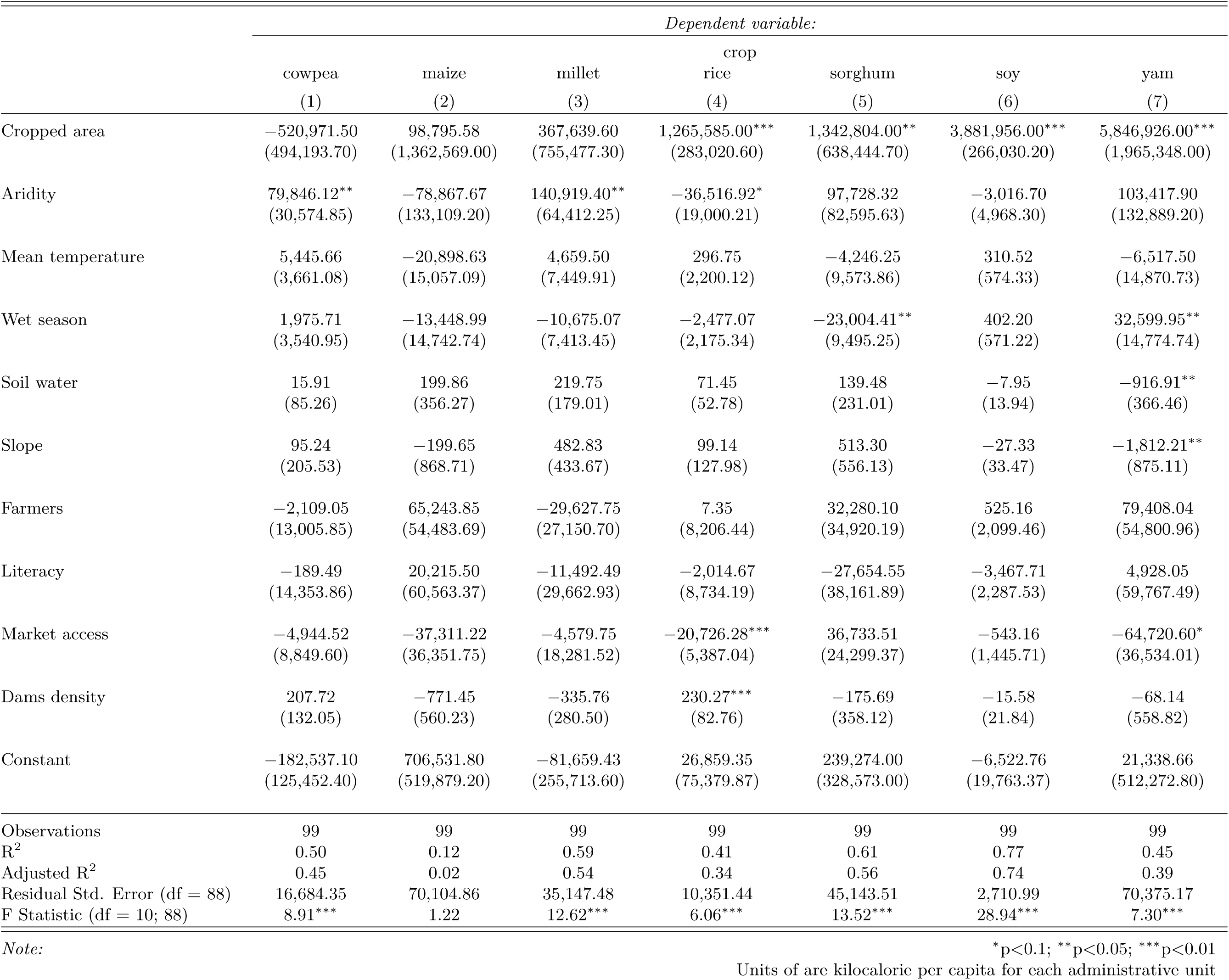
Regression on mean kilocalorie production per crop

|  | *Dependent variable:* |  |  |  |  |  |  |
| --- | --- | --- | --- | --- | --- | --- | --- |
|  | cowpea<br>(1) | maize<br>(2) | millet<br>(3) | rice<br>(4) | sorghum<br>(5) | soy<br>(6) | yam<br>(7) |
| Cropped area | -520,971.50<br>(494,193.70) | 98,795.58<br>(1,362,569.00) | 367,639.60<br>(755,477.30) | 1,265,585.00***<br>(283,020.60) | 1,342,804.00**<br>(638,444.70) | 3,881,956.00***<br>(266,030.20) | 5,846,926.00***<br>(1,965,348.00) |
| Aridity | 79,846.12**<br>(30,574.85) | -78,867.67<br>(133,109.20) | 140,919.40**<br>(64,412.25) | -36,516.92*<br>(19,000.21) | 97,728.32<br>(82,595.63) | -3,016.70<br>(4,968.30) | 103,417.90<br>(132,889.20) |
| Mean temperature | 5,445.66<br>(3,661.08) | -20,898.63<br>(15,057.09) | 4,659.50<br>(7,449.91) | 296.75<br>(2,200.12) | -4,246.25<br>(9,573.86) | 310.52<br>(574.33) | -6,517.50<br>(14,870.73) |
| Wet season | 1,975.71<br>(3,540.95) | -13,448.99<br>(14,742.74) | -10,675.07<br>(7,413.45) | -2,477.07<br>(2,175.34) | -23,004.41**<br>(9,495.25) | 402.20<br>(571.22) | 32,599.95**<br>(14,774.74) |
| Soil water | 15.91<br>(85.26) | 199.86<br>(356.27) | 219.75<br>(179.01) | 71.45<br>(52.78) | 139.48<br>(231.01) | -7.95<br>(13.94) | -916.91**<br>(366.46) |
| Slope | 95.24<br>(205.53) | -199.65<br>(868.71) | 482.83<br>(433.67) | 99.14<br>(127.98) | 513.30<br>(556.13) | -27.33<br>(33.47) | -1,812.21**<br>(875.11) |
| Farmers | -2,109.05<br>(13,005.85) | 65,243.85<br>(54,483.69) | -29,627.75<br>(27,150.70) | 7.35<br>(8,206.44) | 32,280.10<br>(34,920.19) | 525.16<br>(2,099.46) | 79,408.04<br>(54,800.96) |
| Literacy | -189.49<br>(14,353.86) | 20,215.50<br>(60,563.37) | -11,492.49<br>(29,662.93) | -2,014.67<br>(8,734.19) | -27,654.55<br>(38,161.89) | -3,467.71<br>(2,287.53) | 4,928.05<br>(59,767.49) |
| Market access | -4,944.52<br>(8,849.60) | -37,311.22<br>(36,351.75) | -4,579.75<br>(18,281.52) | -20,726.28***<br>(5,387.04) | 36,733.51<br>(24,299.37) | -543.16<br>(1,445.71) | -64,720.60*<br>(36,534.01) |
| Dams density | 207.72<br>(132.05) | -771.45<br>(560.23) | -335.76<br>(280.50) | 230.27***<br>(82.76) | -175.69<br>(358.12) | -15.58<br>(21.84) | -68.14<br>(558.82) |
| Constant | -182,537.10<br>(125,452.40) | 706,531.80<br>(519,879.20) | -81,659.43<br>(255,713.60) | 26,859.35<br>(75,379.87) | 239,274.00<br>(328,573.00) | -6,522.76<br>(19,763.37) | 21,338.66<br>(512,272.80) |
| Observations | 99 | 99 | 99 | 99 | 99 | 99 | 99 |
| R<sup>2</sup> | 0.50 | 0.12 | 0.59 | 0.41 | 0.61 | 0.77 | 0.45 |
| Adjusted R<sup>2</sup> | 0.45 | 0.02 | 0.54 | 0.34 | 0.56 | 0.74 | 0.39 |
| Residual Std. Error (df = 88) | 16,684.35 | 70,104.86 | 35,147.48 | 10,351.44 | 45,143.51 | 2,710.99 | 70,375.17 |
| F Statistic (df = 10; 88) | 8.91*** | 1.22 | 12.62*** | 6.06*** | 13.52*** | 28.94*** | 7.30*** |

Note:
\*p<0.1; \*\*p<0.05; \*\*\*p<0.01 Units of are kilocalorie per capita for each administrative unit

**Table 4:** Regression on kilocalorie variance per crop

|  | *Dependent variable:* |  |  |  |  |  |  |
| --- | --- | --- | --- | --- | --- | --- | --- |
|  | cowpea | maize | millet | rice | sorghum | soy | yam |
|  | (1) | (2) | (3) | (4) | (5) | (6) | (7) |
| Cropped area | -48,361.16<br>(239,633.90) | 821,931.80<br>(514,472.70) | 309,424.50<br>(265,420.70) | 1,375,013.00***<br>(189,838.10) | 1,414,477.00***<br>(177,411.70) | 3,214,423.00***<br>(207,021.20) | 1,455,167.00***<br>(414,967.80) |
| Aridity | 27,280.61*<br>(14,825.71) | -30,600.97<br>(50,258.76) | 35,811.37<br>(22,629.86) | -13,573.50<br>(12,744.53) | 32,330.94<br>(22,951.76) | 1,130.62<br>(3,866.26) | 3,755.60<br>(28,058.50) |
| Mean temperature | 1,435.46<br>(1,775.25) | -6,276.68<br>(5,685.19) | 1,803.32<br>(2,617.36) | 1,550.24<br>(1,475.75) | 206.23<br>(2,660.39) | 301.61<br>(446.94) | 1,689.44<br>(3,139.84) |
| Wet season | -203.79<br>(1,717.00) | -6,507.01<br>(5,566.50) | -3,573.94<br>(2,604.56) | -113.18<br>(1,459.12) | -3,306.13<br>(2,638.55) | 354.40<br>(444.51) | 9,015.52***<br>(3,119.57) |
| Soil water | -10.58<br>(41.34) | -2.98<br>(134.52) | 43.05<br>(62.89) | 7.70<br>(35.40) | 41.96<br>(64.19) | 7.72<br>(10.85) | -114.66<br>(77.37) |
| Slope | 47.54<br>(99.66) | -120.49<br>(328.00) | 164.77<br>(152.36) | -39.67<br>(85.84) | 100.78<br>(154.54) | -23.37<br>(26.04) | -382.35**<br>(184.77) |
| Farmers | 1,699.38<br>(6,306.52) | 21,488.51<br>(20,571.71) | -7,447.55<br>(9,538.81) | -3,605.19<br>(5,504.53) | 6,372.76<br>(9,703.66) | -209.33<br>(1,633.78) | 17,657.47<br>(11,570.79) |
| Literacy | -5,673.88<br>(6,960.17) | 8,255.92<br>(22,867.24) | -8,446.51<br>(10,421.43) | -725.63<br>(5,858.52) | -4,223.14<br>(10,604.47) | -418.62<br>(1,780.12) | -5,567.27<br>(12,619.44) |
| Market access | -20.92<br>(4,291.16) | -10,515.84<br>(13,725.53) | -3,191.11<br>(6,422.82) | -11,694.31***<br>(3,613.39) | 1,949.40<br>(6,752.34) | -1,573.20<br>(1,125.03) | -15,410.34**<br>(7,713.87) |
| Dams density | 80.62<br>(64.03) | -324.33<br>(211.53) | -224.21**<br>(98.55) | 139.11**<br>(55.51) | 6.38<br>(99.51) | -5.51<br>(17.00) | -13.20<br>(117.99) |
| Constant | -41,026.01<br>(60,831.71) | 236,270.00<br>(196,293.60) | -24,970.78<br>(89,839.46) | -28,623.44<br>(50,561.58) | 6,024.16<br>(91,304.21) | -10,020.11<br>(15,379.59) | -87,665.51<br>(108,162.40) |
| Observations | 99 | 99 | 99 | 99 | 99 | 99 | 99 |
| R<sup>2</sup> | 0.49 | 0.13 | 0.56 | 0.52 | 0.71 | 0.79 | 0.52 |
| Adjusted R<sup>2</sup> | 0.43 | 0.03 | 0.51 | 0.47 | 0.67 | 0.76 | 0.46 |
| Residual Std. Error (df = 88) | 8,090.22 | 26,469.88 | 12,348.31 | 6,943.30 | 12,544.53 | 2,109.66 | 14,859.17 |
| F Statistic (df = 10; 88) | 8.33*** | 1.35 | 11.22*** | 9.70*** | 21.04*** | 32.55*** | 9.48*** |

Note:
\*p<0.1; \*\*p<0.05; \*\*\*p<0.01 Units of variance are millions of kilocalorie

**Table 5:** Regression on mean kilocalorie production per crop

|  | *Dependent variable:* |  |  |  |  |  |  |
| --- | --- | --- | --- | --- | --- | --- | --- |
|  | cowpea | maize | millet | crop<br>rice | sorghum | soy | yam |
|  | (1) | (2) | (3) | (4) | (5) | (6) | (7) |
| Cropped area | 338.88**<br>(134.78) | -3.35<br>(23.47) | 106.80<br>(101.72) | 561.04***<br>(157.40) | 161.00**<br>(67.68) | 1,819.15**<br>(716.61) | 249.12<br>(210.94) |
| Aridity | 12.89<br>(8.26) | -2.33<br>(2.26) | 13.44<br>(8.56) | -27.42**<br>(10.47) | 9.39<br>(8.62) | -16.80<br>(13.35) | 2.48<br>(14.13) |
| Mean temperature | 2.33**<br>(0.99) | -0.42<br>(0.26) | 2.86***<br>(0.99) | 1.96<br>(1.21) | 2.18**<br>(1.00) | 1.78<br>(1.54) | -0.81<br>(1.58) |
| Wet season | -0.50<br>(0.96) | 0.06<br>(0.25) | -0.43<br>(0.99) | -2.23*<br>(1.20) | -0.73<br>(0.99) | -0.68<br>(1.54) | 5.48***<br>(1.57) |
| Soil water | 0.04<br>(0.02) | 0.002<br>(0.01) | 0.03<br>(0.02) | 0.08***<br>(0.03) | 0.01<br>(0.02) | 0.05<br>(0.04) | -0.02<br>(0.04) |
| Slope | -0.04<br>(0.06) | -0.003<br>(0.01) | -0.03<br>(0.06) | 0.06<br>(0.07) | 0.03<br>(0.06) | -0.09<br>(0.09) | -0.001<br>(0.09) |
| Farmers | 1.72<br>(3.51) | 0.63<br>(0.93) | 2.97<br>(3.61) | 6.87<br>(4.52) | 5.49<br>(3.64) | 2.28<br>(5.64) | 6.14<br>(5.82) |
| Literacy | -12.68***<br>(3.88) | -1.09<br>(1.03) | -10.75***<br>(3.94) | -6.33<br>(4.81) | -14.55***<br>(3.98) | -10.88*<br>(6.15) | -4.58<br>(6.35) |
| Market access | -7.71***<br>(2.39) | -0.87<br>(0.62) | -8.88***<br>(2.43) | -5.10*<br>(2.97) | -8.21***<br>(2.54) | -12.81***<br>(3.89) | -8.25**<br>(3.88) |
| Dams density | 0.14***<br>(0.04) | -0.003<br>(0.01) | 0.13***<br>(0.04) | 0.06<br>(0.05) | 0.13***<br>(0.04) | 0.06<br>(0.06) | 0.001<br>(0.06) |
| Constant | -49.68<br>(33.87) | 36.59***<br>(8.83) | -65.58*<br>(33.98) | -18.10<br>(41.54) | -41.91<br>(34.28) | -23.90<br>(53.11) | 3.34<br>(54.44) |
| Observations | 99 | 99 | 99 | 99 | 99 | 99 | 99 |
| R<sup>2</sup> | 0.86 | 0.25 | 0.86 | 0.53 | 0.86 | 0.55 | 0.58 |
| Adjusted R<sup>2</sup> | 0.85 | 0.17 | 0.84 | 0.48 | 0.84 | 0.50 | 0.54 |
| Residual Std. Error (df = 88) | 4.51 | 1.19 | 4.67 | 5.70 | 4.71 | 7.29 | 7.48 |
| F Statistic (df = 10; 88) | 55.23*** | 2.97*** | 54.10*** | 10.03*** | 53.55*** | 10.96*** | 12.35*** |

*Note:*
\*p<0.1; \*\*p<0.05; \*\*\*p<0.01
Units of are kilocalorie per capita for each administrative unit in logarithmic scale

**Table 6:**
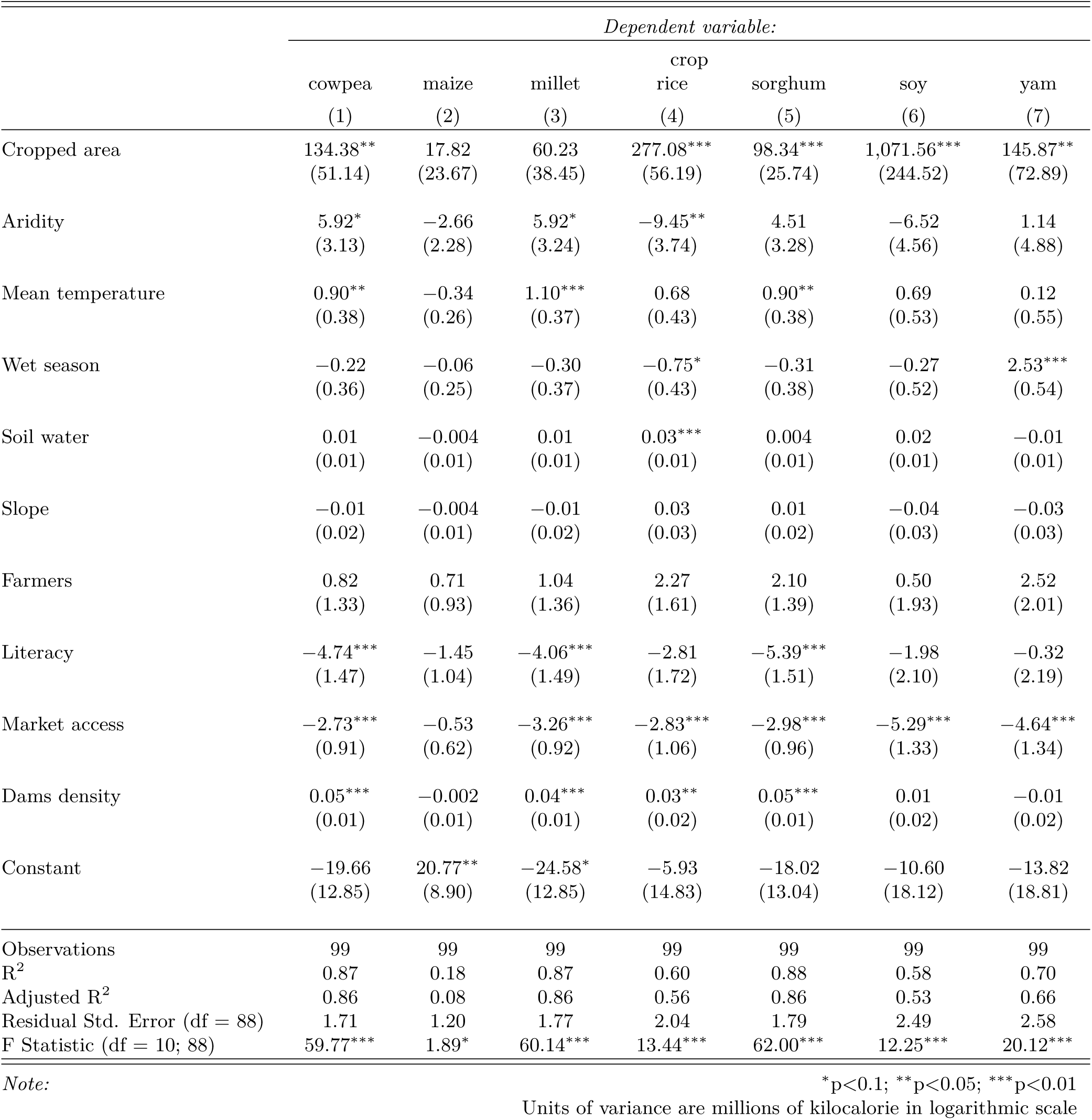
Regression on kilocalorie variance per crop

